# Earliest photic zone niches probed by ancestral microbial rhodopsins

**DOI:** 10.1101/2021.11.18.469010

**Authors:** Cathryn D. Sephus, Evrim Fer, Amanda K. Garcia, Zachary R. Adam, Edward W. Schwieterman, Betül Kaçar

## Abstract

For billions of years, life has continuously adapted to dynamic physical conditions near the Earth’s surface. Fossils and other preserved biosignatures in the paleontological record are the most direct evidence for reconstructing the broad historical contours of this adaptive interplay. However, biosignatures dating to Earth’s earliest history are exceedingly rare. Here, we combine phylogenetic inference of primordial rhodopsin proteins with modeled spectral features of the Precambrian Earth environment to reconstruct the paleobiological history of this essential family of photoactive transmembrane proteins. Our results suggest that ancestral microbial rhodopsins likely acted as light-driven proton pumps and were spectrally tuned toward the absorption of green light, which would have enabled their hosts to occupy depths in a water column or biofilm where UV wavelengths were attenuated. Subsequent diversification of rhodopsin functions and peak absorption frequencies was enabled by the expansion of surface ecological niches induced by the accumulation of atmospheric oxygen. Inferred ancestors retain distinct associations between extant functions and peak absorption frequencies. Our findings suggest that novel information encoded by biomolecules can be used as “paleosensors” for conditions of ancient, inhabited niches of host organisms not represented elsewhere in the paleontological record. The coupling of functional diversification and spectral tuning of this taxonomically diverse protein family underscores the utility of rhodopsins as universal testbeds for inferring remotely detectable biosignatures on inhabited planetary bodies.

## INTRODUCTION

Decoding the complex relationships between life and the environments it inhabits is central to reconstructing the factors that determine planetary habitability over geologic timescales. Similarly, this approach is necessary to constrain the physical limits of biological functionality, identify general principles that may govern the evolution of living systems, and aid the search for life elsewhere in our universe (Harrison 2013; McKay 2014). Planetary habitability is strongly influenced by solar insolation and photon radiance flux, which couple planetary and stellar evolution to the persistence of life (Newman 1977; Lunine 2006). Much of Earth’s historical geochemical proxy record is incomplete, which imposes significant limitations on the reconstruction of planetary and solar factors that have influenced habitability and the co-evolution of life and its environments.

The information encoded in life itself may provide novel insights into how our planet has maintained planetary habitability where geologic and stellar inferences fall short. The earliest paleontological record lacks abundant, direct evidence of major metabolic and morphogenetic features (Butterfield 2007; Garcia et al. 2021), but the contemporary bioinformatic record is incredibly rich and includes several environment-sensitive biomolecular components. Reconstructing the ancestral sequences of biomolecules that are strongly coupled to habitability parameters may provide a means of extrapolating ancient biological phenotypes—and the environmental conditions that shaped them—back to the Last Universal Common Ancestor (LUCA) (Doolittle 2000; Benner 2002; Ruiz-Gonzalez and Marin 2004; Garcia 2019).

Photosensitive proteins are key intermediaries that connect intracellular chemical states, extracellular substrate availability, and solar irradiance. These biomolecules constitute a promising system for tracking ancient physical parameters that are not directly recorded in the geologic record. All known phototrophic metabolisms on Earth rely on one of three energy-converting pigments which transform light energy into chemical energy. These pigments include chlorophylls, bacteriochlorophylls, and retinal (Gómez-Conarnau 2019). Retinal-based pigment proteins, known as rhodopsins, have been found in Archaea, Bacteria, Eukarya, and giant viruses (Bryant and Frigaard 2006; Pinhassi 2016). Absolute age constraints on the oldest rhodopsins are unknown and the subject of investigation. Rhodopsin distribution across such diverse taxa has been attributed to high rates of microbial gene transfer and not necessarily evidence of rhodopsin presence in the last universal common ancestor (LUCA) (Sharma 2006). Other analyses suggest that the taxonomic distribution of rhodopsins may still be indicative of ancient origins, possibly predating the divergence of the three domains of life (Ruiz-Gonzalez and Marin 2004). Finally, it has been proposed that a primitive ion translocation functionality, perhaps comparable to that carried out by extant microbial rhodopsins, may have been present in some of the most deeply-rooted ancestors of today’s microorganisms (Shalaeva 2015).

Retinal-based phototrophy utilizes a light-capture system that is simpler than its chlorophyll-based counterparts (DasSarma and Schwieterman 2018). Rhodopsin pigments are composed of one seven-transmembrane opsin protein and one retinal chromophore (Mackin 2014). Photoisomerization of the retinal chromophore within the opsin active site (also referred to as the retinal binding pocket) triggers a series of cyclical conformational changes that can directly drive ion transport across the cellular membrane, generating a proton-motive force that may be harnessed by ATP synthase to generate ATP (Racker 1974). The simple bipartite architecture of rhodopsins has also been harnessed by biology for uses beyond ion transport, including both phototaxis and photophobia, substrate uptake, and starvation prevention (Sharma 2006; Steindler 2011; Akram 2013; Govorunova 2017). Rhodopsins can be tuned to absorb photons across much of the visible light spectrum (Engqvist et al. 2015) and have been sampled from exposed terrestrial surfaces to the sub-photic marine zone (Inoue et al. 2018). Today, microbial (type 1) rhodopsins are among the dominant energy-transducing agents that harvest solar energy in the surface ocean (Gómez-Conarnau 2019). It was suggested that the structural simplicity and functional diversity of microbial (type 1) rhodopsins may have promoted their spread to global econiches (Schwieterman et al. 2018). The combination of taxonomic distribution, phylogenetic tractability, and niche ubiquity indicates that ancestral type 1 rhodopsins are excellent candidate molecular systems for characterizing long-term conditions of habitability and paleoecology where geologic indicators are unavailable (Finkel et al. 2012; Kacar et al. 2017; Lamsdell et al. 2017).

We reconstructed the evolutionary history of microbial type 1 rhodopsins to explore ancestral properties that might serve as a guide for understanding early Earth paleoecology and habitability. Protein sequence analyses of both extant and phylogenetically inferred ancestral rhodopsins were used to constrain possible ancestral rhodopsin functions. In addition, maximum absorbance wavelength of ancestral rhodopsins was predicted by a machine-learning algorithm that models the relationship between spectral absorbance peaks and amino acid sequence (Karasuyama et al. 2018). Our analyses suggest proton-pump functionality and green wavelength absorption for the last common ancestor of ancestral type 1 rhodopsins. Green wavelength absorption is parsimonious with our modeled spectral features of the Precambrian Earth environment, providing an adaptive advantage to shallow coastal marine niches with abundant nutrients, metabolically useful light availability and attenuated phototoxic UV irradiation. The proliferation of specialized functions and the spectral tuning of rhodopsins likely resulted from the ecological upheaval wrought by the rise of atmospheric oxygen, with significant accumulation marked by the interval known as the Great Oxidation Event (GOE) approximately 2.3 billion years ago (Lyons et al. 2014; Sánchez-Baracaldo et al. 2021). The convergence of our results from reconstructed, ancestral biomolecular characteristics and indirect physical models reveals the manner by which phototrophic organisms have adapted to changing habitability parameters over billions of years.

## RESULTS AND DISCUSSION

### Conservation and statistical support for ancestral rhodopsin residues important for function and spectral tuning

We assembled a dataset of representative protein sequences with which to reconstruct the evolutionary history and ancestral traits of microbial rhodopsins. BLASTp (Camacho et al. 2009) was used to identify 452 homologous type 1 rhodopsin sequences from the National Center for Biotechnology Information (NCBI) non-redundant and UniProt protein databases. We additionally identified 116 homologs belonging to heliorhodopsins, a recently discovered rhodopsin class that exhibits an inverted membrane topology relative to type 1 rhodopsins and constitutes a distinct outgroup (Pushkarev 2018; Lu et al. 2019; Rozenberg et al. 2021). Animal, or type 2, rhodopsins, though exhibiting a similar seven transmembrane architecture, possess little sequence homology with microbial type 1 rhodopsins and were thus excluded from the dataset (Sharma 2006; Rozenberg et al. 2021). Our type 1 rhodopsin sequence dataset comprises four major archaeal rhodopsin groups: 1) bacteriorhodopsin, outward proton pumps; 2) halorhodopsin, inward chloride pumps; 3) sensory rhodopsin I, positive phototaxis mediators; and 4) sensory rhodopsin II, negative phototaxis mediators (Ernst 2014). The dataset also includes two major bacterial rhodopsin groups: 1) proteorhodopsin and 2) xanthorhodopsin, both outward proton pumps, with the latter distinguished by the ability to bind a second carotenoid chromophore (Ernst 2014). These taxonomically and functionally distinct lineages were resolved into their respective clades by maximum likelihood phylogenetic reconstruction (Fig. 1), producing a tree topology largely in agreement with previous analyses (Ihara 1999; Pinhassi 2016; Shibata 2018; Rozenberg et al. 2021). Alternate phylogenetic trees, generated by incorporating next-best-fit evolutionary models, reproduced this general topology (see Methods; Supplementary Fig. 1). Ancestral amino acid sequences were inferred by maximum likelihood from internal nodes of our rhodopsin phylogeny. Five of the oldest internal nodes were selected as targets for further phenotypic analysis: 1) Anc1, ancestral to bacteriorhodopsins and halorhodopsins, 2) Anc2, ancestral to sensory rhodopsins, 3) Anc3, ancestral to archaeal rhodopsins, 4) Anc4, ancestral to bacterial rhodopsins, and 5) Anc5, ancestral to all type 1 rhodopsins (Fig. 1, Supplementary Fig. 2).

**Fig. 1.**
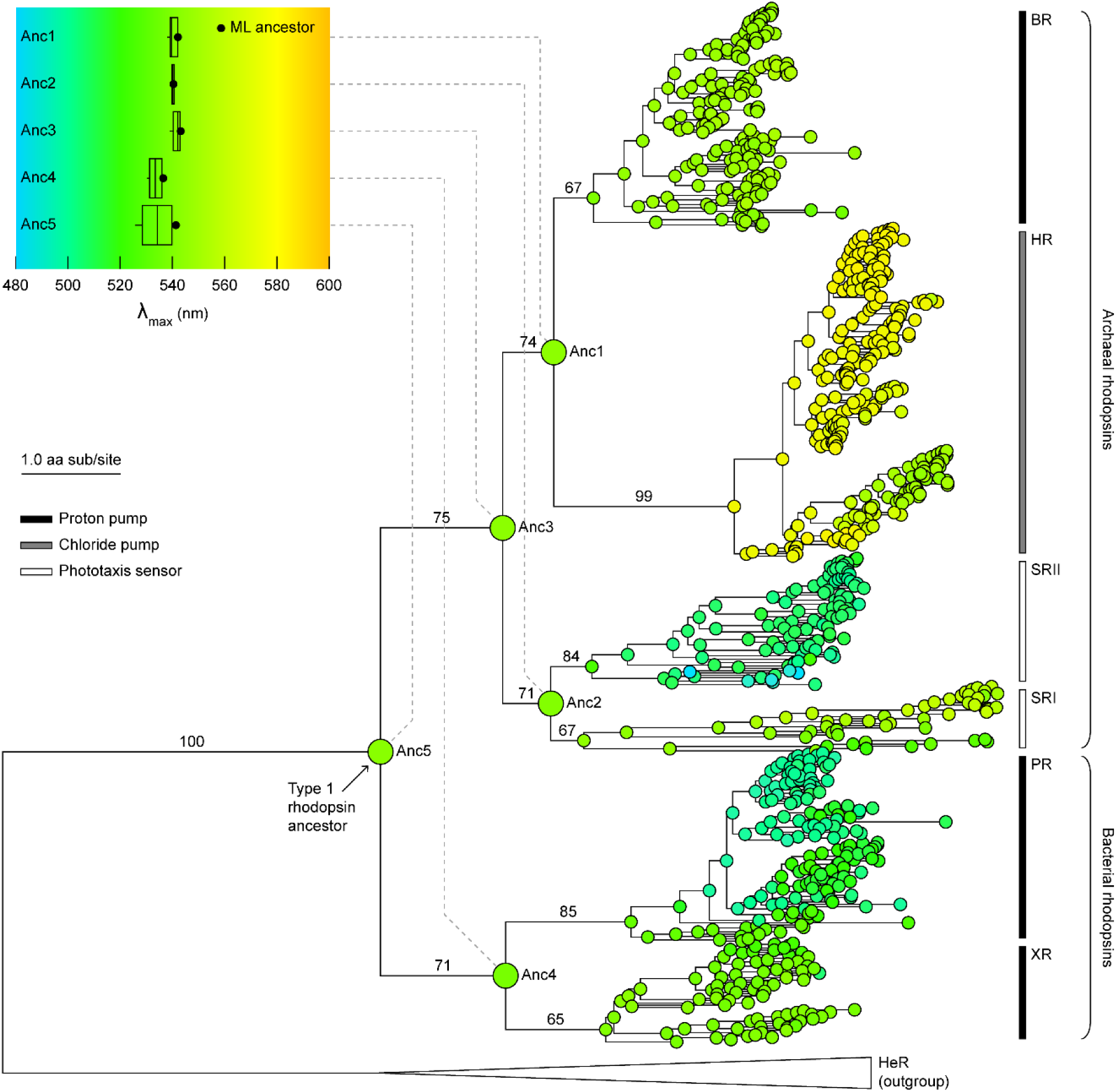
Evolution of spectral tuning mapped to a maximum likelihood phylogeny of microbial type 1 rhodopsin protein sequences. Nodes are colored according to the predicted maximum absorption wavelength (λ_max_) of the associated maximum likelihood (ML) ancestral sequence (color scale shown in plot, top left). Boxplot shows the predicted λ_max_ distribution among alternate ancestral sequences for nodes Anc1-Anc5 (n = 50 per node; see Methods and Fig. 2b,c). Branch support values are derived from 100 bootstrap replicates. Branch scale indicates 1.0 amino acid substitutions per site. Heliorhodopsin outgroup clade is collapsed. BR: bacteriorhodopsin; HR, halorhodopsin; SR II, sensory rhodopsin II; SR I, sensory rhodopsin I; PR, proteorhodopsin; XR, xanthorhodopsin; HeR, heliorhodopsin.

Due to their deep position in the phylogenetic tree, mean site posterior probabilities for ancestors Anc1-Anc5 range between 0.55 and 0.76 (Fig. 2a; Supplementary Table 1). However, statistical support is enriched for 23 ancestral binding pocket residues (<5 Å from the retinal chromophore), constituting the most conserved element of microbial rhodopsin structure (Kandori 2015). Mean posterior probabilities for binding pocket residues fall between 0.90 and 0.97 (Supplementary Table 1, 2). Nine of these residues have ≥95% mean pairwise identity across all extant sequences included in our dataset (Y83, W86, P91, G122, W182, Y185, W189, D212, and K216 in the bacteriorhodopsin of *Halobacterium salinarum*). Most have significant, experimentally determined contributions to rhodopsin function (Greenhalgh 1993; Kusnetzow 1999; Babitzki 2009; Kandori 2015; Ding 2018), including the strictly conserved K216 residue that is covalently bonded to retinal via a Schiff base linkage (Babitzki 2009). Other functions include maintenance of the internal charge environment for ion-pumping rhodopsins (Song and Gunner 2014; Kandori 2015) or mediation of transducer interactions required for phototactic response in sensory rhodopsins (Sudo et al. 2006). Finally, binding pocket residues have been implicated as having outsized influence in rhodopsin color tuning (Ren et al. 2001; Man 2003; Inoue 2021). Thus, the enriched statistical support for ancestral binding pocket residues permits sequence-based inferences of historical rhodopsin functionality.

**Fig. 2.**
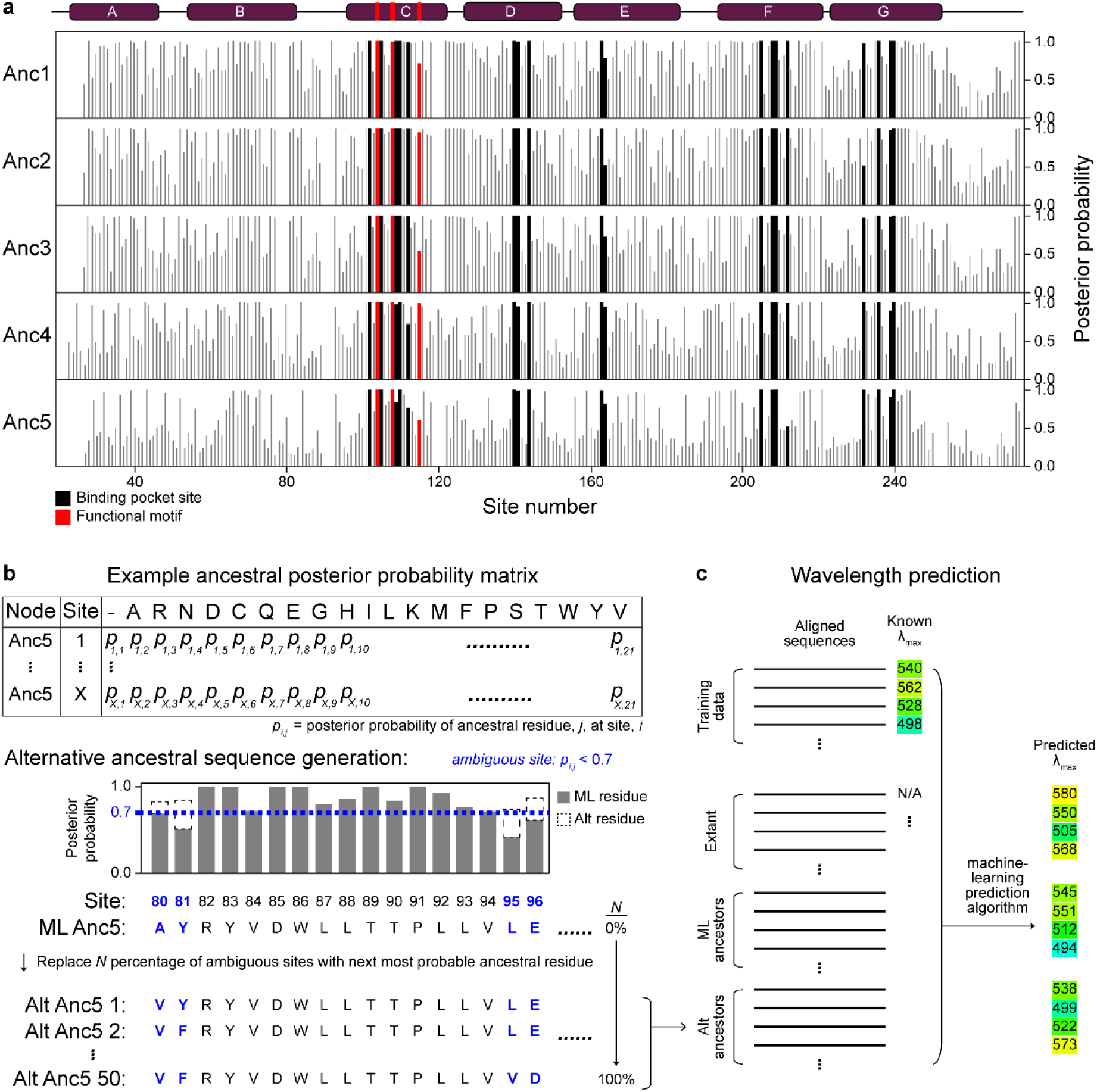
Incorporation of statistical uncertainty of ancestral rhodopsins inference into wavelength prediction. (*A*) Site-wise posterior probabilities of maximum likelihood (ML) ancestral rhodopsin sequences. Sites labeled “functional motif” are homologous to the three-residue DTD motif in extant *H. salinarum* bacteriorhodopsin. Helices are labeled above. (*B*) Generation of alternative (“alt”) ancestral sequences. For each node, alternate sequences were generated by replacing 0% to 100% of ambiguous sites (defined as having a posterior probability < 0.7) with the next most probable residue. (*C*) Methodology for maximum absorbance wavelength (λ_max_) prediction. λ_max_ values of extant and ancestral (both ML and alternative) rhodopsin sequences were predicted by a machine-learning model (group-LASSO-based machine learning method), trained on a dataset of experimentally characterized rhodopsin proteins with known λ_max_ values.

### Ancestral type 1 rhodopsins likely acted as light-driven proton pumps

Comparative analysis of extant variants has provided compelling circumstantial evidence that primordial rhodopsins functioned as transporters (Ihara 1999), but the evolutionary progression of functions or of photon absorptivity have not been explicitly demonstrated to date. As an initial, global approach to constraining early rhodopsin functionality, 3D protein structures of maximum likelihood ancestral sequences Anc1-Anc5 were predicted with Phyre2 (Kelley et al. 2015) and used for functional annotation by the COFACTOR algorithm within the I-TASSER software pipeline (Roy 2010). Modeled ancestral rhodopsin structures preserve the characteristic seven-transmembrane alpha-helical fold, otherwise known as the G-protein coupled receptor fold (Mackin 2014)—of extant microbial type 1 homologs (Fig. 3). Kyte and Doolittle hydropathy plots for Anc1-Anc5 reveal seven highly hydrophobic regions, consistent with the conservation of seven transmembrane domains (Supplementary Fig. 3). Membrane localization of ancestral rhodopsins was also predicted by the SOSUI tool (Hirokawa et al. 1998) (Supplementary Table 3). Additionally, the ∼21-residue-long C-terminal sequence of these ancestral rhodopsins form a distinct hydrophilic region, indicating an orientation towards the cytoplasm as observed with extant type 1 rhodopsins (Pushkarev 2018). Finally, the structure-based functional predictions are consistent with early type 1 rhodopsins that carried out phototransduction, formed protein-chromophore linkages, and acted as proton transmembrane transporters (Supplementary Table 4).

**Fig. 3.**
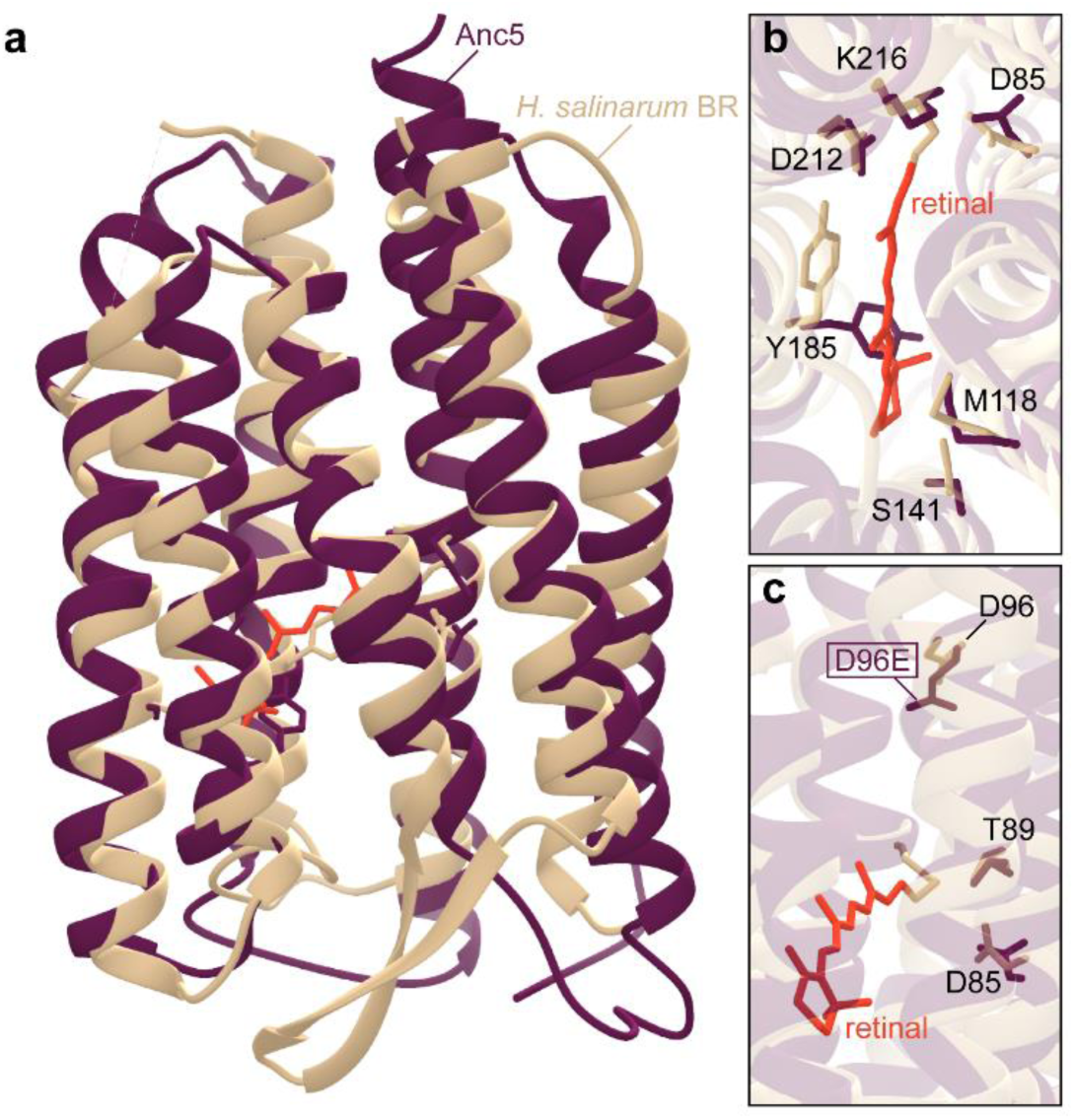
Structural conservation between the inferred, type 1 rhodopsin last common ancestor, Anc5, and extant bacteriorhodopsin. (*A*) Homology model of Anc5 aligned to the extant *H. salinarum* bacteriorhodopsin (“BR”; PDB: 1C3W). (*B*) Retinal binding pocket of ancestral and extant rhodopsins. Conserved residues that interact with the retinal chromophore are labeled. (*C*) Residues homologous to the *H. salinarum* DTD motif correlative with rhodopsin function. Extant bacteriorhodopsin exhibits a DTD motif, compared to the DTE motif in Anc5. (*B,C*) All displayed Anc5 residues are identical to those of extant bacteriorhodopsin except for site 96 (D96E, boxed).

Given the comparatively high statistical support and functional significance of ancestral binding pocket residues, we interrogated these specific sequence features to provide complementary insights into plausible functionality of ancient rhodopsins. For extant, uncharacterized rhodopsins, functional annotation is often guided by a characteristic, three-residue motif that includes sites homologous to the D85, T89, and D96 residues (a.k.a., “DTD” motif) in bacteriorhodopsin (*H. salinarum*) (Kandori 2015; Hasegawa et al. 2020; Inoue 2020; Rozenberg et al. 2021). These sites are located within a section of the third transmembrane helix (“Helix C”; Fig. 3c, Fig. 4) that constitutes a portion of the retinal binding pocket. Variation in this motif is strongly correlated with functional differentiation in rhodopsins. For example, “DTD” or “DTE” motifs (glutamic acid replaces the D96 residue in the latter) are characteristic of bacteriorhodopsins, xanthorhodopsins, and proteorhodopsins, which all function as light-driven proton pumps and exhibit similar photosensitive mechanisms (Kandori 2015). By contrast, chloride-pumping halorhodopsins exhibit a “TSA” motif and sensory rhodopsins instead exhibit a “DTY” or “DTF” motif (Engelhard et al. 2018).

**Fig. 4.**
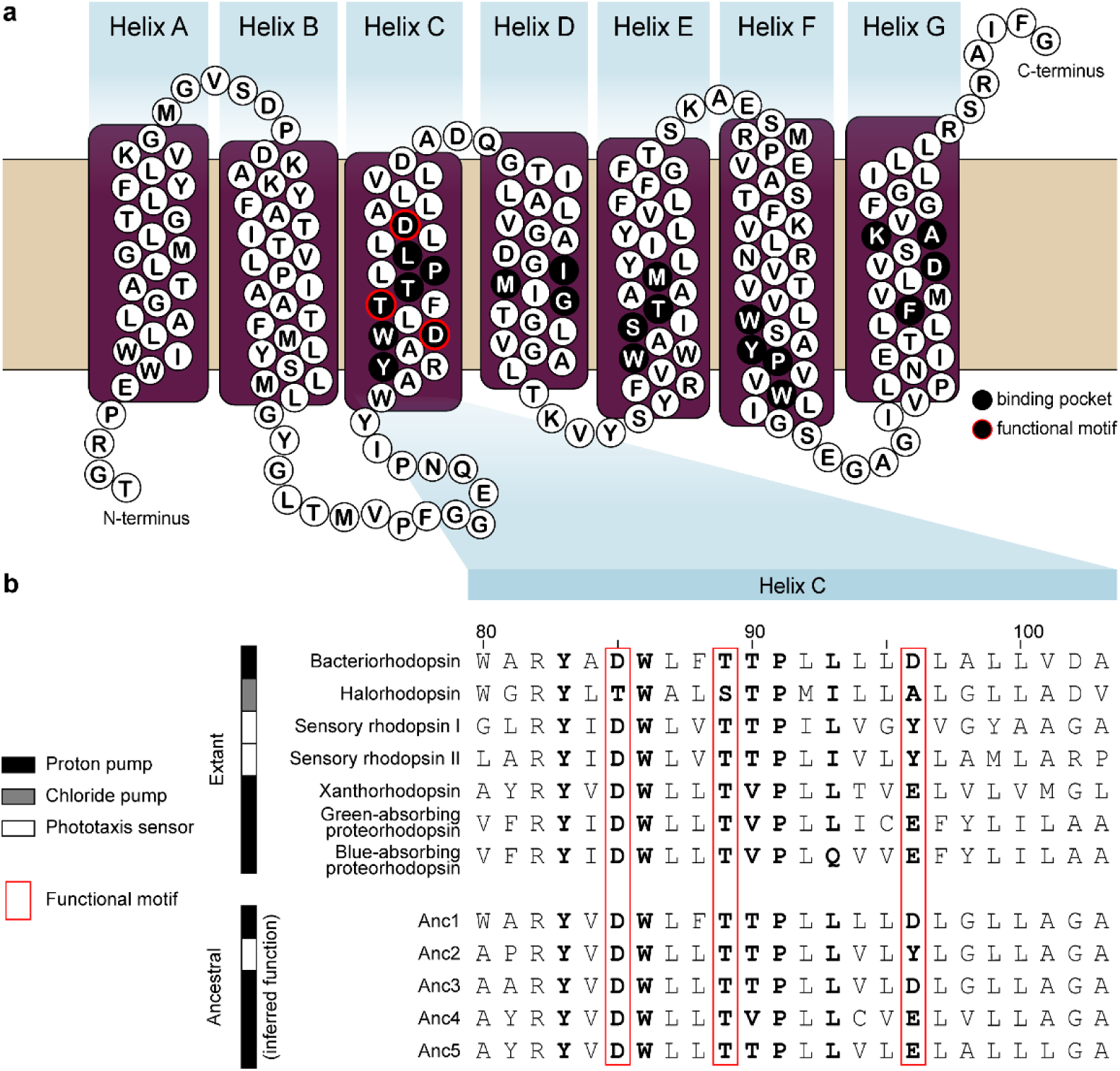
Rhodopsin binding pocket residues in extant and inferred ancestral sequences. (*A*) Secondary structure of extant *H. salinarum* bacteriorhodopsin, indicating the binding pocket residues as well as the functionally correlative DTD motif. (*B*) Protein sequence alignment of Helix C residues for representative, extant type 1 rhodopsins and ancestral rhodopsins. Binding pocket residues are indicated by bold font. (*A,B*) Sites labeled “functional motif” are homologous to the three-residue DTD motif in extant *H. salinarum* bacteriorhodopsin.

The functional correlation of the three-residue motif can be explained by the role these sites play during ion binding and transfer. In proton-pumping rhodopsins, the negatively charged residues of the DTD/DTE motifs are protonated during intermediate steps of the bacteriorhodopsin, xanthorhodopsin, and proteorhodopsin photocycles, permitting unidirectional proton transport out of the cytoplasm (Kandori 2015). By contrast, the TSA motif typical of chloride-pumping rhodopsins lacks a negative charge relative to the DTD/DTE motifs, which is compensated by binding of the chloride ion (Besaw et al. 2020; Inoue 2020). The functional significance of these motifs has been underscored by experiments demonstrating the conversion of proton-pumping bacteriorhodopsins into chloride pumps by a D85T mutation at the first site of the DTD motif (Sasaki et al. 1995). In certain cases, functional predictions of extant rhodopsins based on these characteristic motifs have been shown to prove reliable upon experimental characterization (Hasegawa et al. 2020).

At the deepest nodes in the rhodopsin phylogeny, the most probable ancestral motifs are: 1) Anc1, DTD, 2) Anc2, DTY, 3) Anc3, DTD, 4) Anc4, DTE, and 5) Anc5, DTE (Fig. 3). These motifs suggest a proton-pumping function for all ancestors bearing the DTD/DTE motifs, except for Anc2, which, as the common ancestor to sensory rhodopsins I and II, exhibits the DTY motif consistent with extant sensory rhodopsins. Most inferred residues at these three sites have posterior probabilities >0.90 for Anc1-Anc5 (Supplementary Table 2). The exceptions are in Anc1, Anc3, and Anc5, for which the third site in the motif has a posterior probability of 0.71, 0.54, or 0.61, respectively. However, the next most likely ancestral states for these sites are either aspartic acid or glutamic acid, maintaining the protonated carboxylic group observed in extant proton transporters. Thus, a motif consistent with an ancestral proton-pumping function is well supported for Anc1, Anc3, Anc4, and Anc5, which together suggests that other microbial functions such as other forms of ion transport and phototaxis may be derived from an ancestral, light-driven proton pump functionality. This model is compatible with experimental findings that sensory rhodopsins, when expressed in alternate microbial hosts in the absence of associated transducer proteins, retain the ability to transport protons, suggesting that the photosensory function is more derived (Bogomolni et al. 1994).

The early evolution of proton transport by rhodopsins or rhodopsin-like proteins would likely have provided an adaptive, simplistic means of maintaining the intracellular environment of ancient cells. Life on early Earth likely faced several hurdles related to membrane permeability that implicate the presence of relatively simple, membrane-spanning motive proteins proximal to LUCA (Pohorille et al. 2003; Deamer and Dworkin 2005; Mulkidjanian et al. 2009). One of these hurdles was circumventing the disruptive cellular tendency to expand and lyse due to the influx of salts from their environment passing across their semi-permeable protocell membrane (Pohorille 2009). The simplistic membrane-spanning ion-pump infrastructure of an early rhodopsin could have provided cells inhabiting photic environments with the means to counteract diffusive flux with a mechanism of active transport (Maloney 1985; Damer 2020). Though the age of earliest proton-pumping rhodopsins is not constrained, biological preference for transporting cations in lieu of anions is consistent with the proposed acidic conditions of the early Archaean ocean, where a high concentration of chloride-based salts such as H^+^, Na^+^, K^+^, Ca^+^, and Mg^+^ would have been prevalent (Damer 2020; Ueda 2021). Acquiring a means of agency over intercellular ion gradient dynamics would have permitted the maintenance of transmembrane potential and chemiosmotic coupling for energy generation, a property which may have been present in LUCA (Lane et al. 2010; Penny et al. 2014; Shalaeva 2015). From a bioenergetic perspective, our results support the premise that a rhodopsin-like proton pump could have enabled chemiosmotic coupling and simplistic energy production in primordial cells (Racker 1974; Berhanu et al. 2019).

### The type 1 rhodopsin ancestor was likely tuned to absorb green light

Extant rhodopsin variants exhibit absorption capacity across much of the visible spectrum. It is unclear which portion of the spectrum the earliest rhodopsins would have attenuated, nor whether changes in absorption would have been correlated with the diversification of new functions. To understand the evolution of rhodopsin spectral tuning, we used a recently developed group-LASSO-based machine-learning method (Karasuyama et al. 2018). This method approximates the maximum absorbance wavelength (λ_max_) of rhodopsin protein sequences by producing a statistical model that describes the relationship between amino acids within the seven transmembrane helices and absorbance features of experimentally characterized homologs. The model was demonstrated to predict λ_max_ of rhodopsins with an average error of ±7.8 nm. Here, λ_max_ values were predicted for extant sequences in the phylogenetic dataset, inferred maximum likelihood ancestors, and alternative ancestors for nodes Anc1-An5 generated to test the robustness of wavelength prediction to the statistical uncertainty of ancestral sequence inference (see Methods; Fig. 2b,c)

The first-order spectral patterns predicted for ancestral rhodopsins indicate lineage- and, potentially, function-specific color tuning. λ_max_ for all ancestors range between ∼480 to 585 nm, occupying the cyan-to-yellow range of the visible light spectrum (Fig. 1). Different rhodopsin lineages, in addition to being associated with varying functions, also exhibit characteristic λ_max_ values. For example, our results suggest that halorhodopsins evolved to absorb yellow light (∼565-585 nm) early, with secondary shifts toward green light absorption in a specific sublineage. The bacteriorhodopsin, sensory rhodopsin I, and xanthorhodopsin clades largely exhibit green spectral tuning (∼500-565 nm), whereas shifts towards cyan (∼480-500 nm) absorption can be observed within the sensory rhodopsin II and proteorhodopsin clades. Notable are the so-called blue-light and green-light absorbing members of the proteorhodopsin clade, whose differential spectral tuning has been suggested to correlate with habitat depth of host microbes in the marine environment (Bielawski et al. 2004; Ozaki et al. 2014). Because green light is attenuated at shallower depths in clear water with minimal particulate matter, it has been argued that blue-absorbing rhodopsins are hosted by organisms that occupy comparatively deeper portions of the water column. Our λ_max_ predictions suggest that blue-light absorbing proteorhodopsins are derived from a green-light absorbing ancestor. This spectral property would have been adaptive for a shallow marine environment where blue light is preferentially absorbed or scattered by suspended inorganic and organic material and the transmission of green light is favored (Mascarenhas and Keck 2018).

All oldest, maximum likelihood ancestral sequences (Anc1-Anc5) from our rhodopsin phylogeny have predicted λ_max_ values between 537 and 543 nm (Fig. 1). Anc5 is the deepest ancestral node, having both the greatest ancestral sequence uncertainty and, by extension, the greatest variation (∼20 nm range) in predicted λ_max_ values across alternate ancestral reconstructions. Accounting for alternate ancestral sequence reconstructions at these same nodes, the predicted λ_max_ range for early ancestors is only expanded to ∼525 to 545 nm. This range is still well within the green portion of the visible light spectrum, even if additionally accounting for the ±7.8 nm average error associated with the wavelength prediction algorithm (Karasuyama et al. 2018). In addition, these results are robust to the size and composition of the training dataset, yielding a comparable ∼524 to 551 nm range across different subsampling methods (**Supplementary Table 5**). Together, our spectral tuning analyses support a model wherein extant type 1 rhodopsins diversified from an ancestor tuned toward the absorption of green light.

### Indicators of early rhodopsin host niche habitability and coevolution with Earth environment

Together, our results indicate that ancestral rhodopsins transported protons across the cellular membrane and exhibited spectral tuning toward green photons over a span of time encompassing Earth’s early history of inhabitation. These inferences are robust both to the statistical uncertainty of our ancestral sequence reconstructions as well as the average error associated with the implemented machine-learning algorithm for predicting ancestral rhodopsin absorbance features. Despite the inherent limitations of computational analyses, our approach provides plausible constraints on early rhodopsin properties across hundreds of ancestral sequences of diverse taxonomic and functional groups. Such predictions can be further tested by the laboratory reconstruction and characterization of ancestral rhodopsins reported here.

Correlations between tuning of photon absorption and cellular function across modern variants might present a logical basis to infer the most likely niche that hosts of primordial, microbial rhodopsins would have occupied. Paleoecological and paleoenvironmental inferences based on organism-level associations and anatomical attributes are well-established in paleobiology, particularly for ecosystems that existed billions of years ago and for which sedimentological indicators are poor, absent or contested (Hofmann and Jackson 1994; Butterfield and Rainbird 1998; Adam et al. 2017). Inferences of ‘phylogenetic paleoecology’ based on molecule-level attributes constitute a relatively new approach to these problems (Kacar et al. 2017; Lamsdell et al. 2017).

Color tuning over microbial rhodopsin evolution has likely been shaped by a complex interplay of both functional and environmental constraints (Man 2003; Bielawski et al. 2004; Engqvist et al. 2015; DasSarma and Schwieterman 2018; Tsujimura and Ishikita 2020). Regarding the former, it is not unexpected that both spectral absorbance patterns and protein function are evolutionarily correlated, given the observation that many of the same rhodopsin binding pocket sites strongly determine both features (Engqvist et al. 2015; Karasuyama et al. 2018; Tsujimura and Ishikita 2020). Our analysis likewise indicates that λ_max_ values correlate with cellular function throughout rhodopsin’s history. The oldest common ancestors of each clade possess comparable λ_max_ values to modern counterparts, which themselves appear generally binned according to functionally distinct lineages (Fig. 1).

Spectral tuning may be further segregated by the photon availability of the host environment and by selective pressures to enable simultaneous optical control of distinct functions by different colors of light. Host organisms containing rhodopsin variants are found almost everywhere near Earth’s surface, occupying a variety of different niches. For example, chloride-pumping halorhodopsins are mostly limited to halophilic Archaea (Ugalde et al. 2011) and maximally absorb yellow light, matching the spectral insolation peak experienced in the terrestrial salt ponds in which they inhabit. By contrast, sensory rhodopsin II and proteorhodopsin absorbances hosted in shallow-water photic zones are shifted towards blue light, where peak spectral irradiance is attenuated by water transmissivity. Sensory rhodopsin I (SRI) tuning is unique because it appears to have been shaped both by its dual roles in phototactic and photophobic responses, as well as by the absorption profiles of other rhodopsin proteins in the same host. In a host halobacterium, orange light activates SRI, which directs flagellar motion towards the light source, but absorption of extra photons in the UV triggers another cascade that stops flagella motion to avoid phototoxicity (Spudich and Bogomolni 1984; Wilde and Mullineaux 2017), thereby guiding the host to wavelengths where both halorhodopsin and bacteriorhodopsin absorb maximally (Sharma 2006).

Inferring the environmental constraints on early rhodopsin evolution is complicated by the fact that Earth’s atmospheric evolution has drastically altered the surface photon irradiance profile over billions of years. Thus, we reconstructed photon irradiance profiles at various water depths for both modern and haze-free Archean Earth scenarios, informed by an updated prescription for the composition of Earth’s early atmosphere with high carbon dioxide and methane, but no oxygen (Arney 2016) (Fig. 5). These models coupled an advanced atmospheric radiative transfer model SMART (Meadows 1996) and the absorption profile of clear liquid water (Segelstein 1981). We found that the level of shielding enjoyed at the surface of Earth today in a haze-free Archean atmosphere is only possible at depths of tens of meters or deeper, and that the equivalent protective depth level is a strong function of wavelength. At visible wavelengths, red and infrared photons also become strongly attenuated with depth, leading to a wavelength of maximum transparency at ∼485 nm in the blue-green portion of the visible spectrum (assuming no impurities, particulate matter will preferentially scatter blue light over green light). This contrasts with a peak spectral irradiance at Earth’s surface without water attenuation, which ranges from roughly 500-530 nm with a corresponding color of cyan-green.

**Fig. 5.**
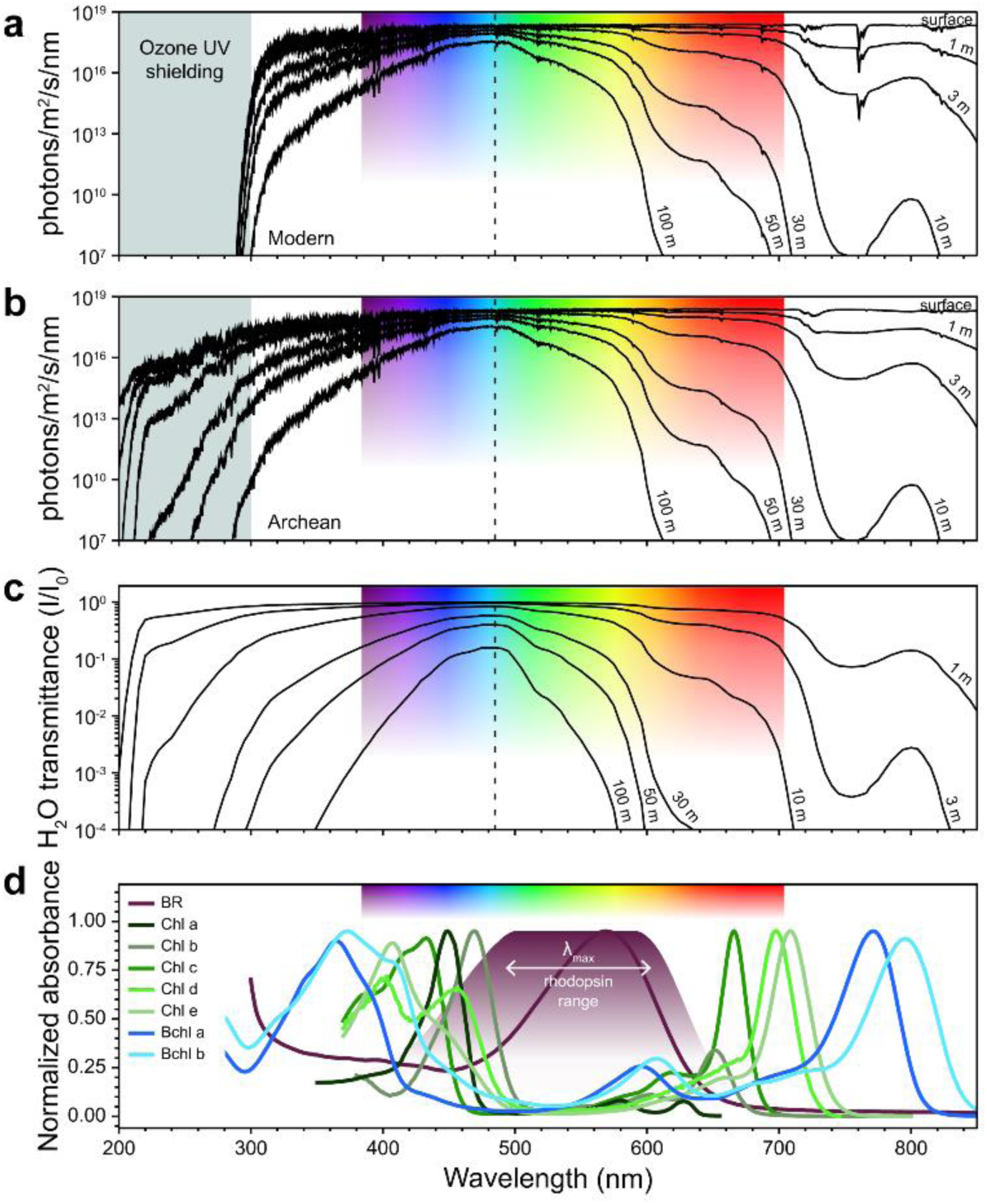
Diversity of biological pigment absorbance profiles in the context of changing surface photon irradiance over Earth history. (*A,B*) Photon irradiance profiles at varying water depths, given a modern (*A*) or Archean (*B*) atmospheric composition. (*C*) Water transmission profile at varying water depths. Transmission was calculated from the coefficients of Segelstein (1981) and the Beer-Lambert Law. (*A,B,C*) Vertical dashed line indicates the maximum water transmission wavelength of 485 nm. (*D*) Normalized absorbance for bacteriorhodopsin (“BR”), chlorophylls (“Chl”), and bacteriochlorophylls (“Bchl”). Shaded purple region indicates the extant and ancestral range of predicted rhodopsin absorption peaks in Fig. 1. Chlorophyll and bacteriochlorophyll spectra are sourced from the Virtual Planetary Laboratory’s Biological Pigments Database (http://vplapps.astro.washington.edu/pigments), with contributions from Jeffrey (1963), Frigaard (1996), and Chen (2010, 2011). The bacteriohodopsin spectrum was sourced from DasSarma and Schwieterman (2018).

In addition to reconstructing historical changes in Earth surface irradiation, we explored the impact that the GOE (Lyons et al. 2014), one of the most significant environmental transitions in Earth-life coevolution, may have had in the evolutionary diversification of microbial type 1 rhodopsins. Light and oxygen both destroy the rhodopsin-associated retinal easily (Hampp and Oesterhelt 2004), so the various functions of rhodopsins as sensors of these external parameters are intimately tied to sequence changes throughout the phylogenetic family. The last common ancestor of haloarchaea is thought to have possessed the halorhodopsin (and perhaps a bacteriorhodopsin) gene (Sharma 2007). As haloarchaea are oxygen-consuming heterotrophs, they would likely have originated only after the rise of oxygenic photosynthesis (Schwieterman et al. 2018), constraining the maximum age of the halorhodopsin clade to at least the GOE at ∼2.3 Ga, and perhaps as far back as the first indicators of oxygenation around 3.0 Ga (Olson et al. 2013; Cardona 2018). Furthermore, sensory rhodopsin II further drives a photophobic response under conditions of abundant oxygen to avoid photooxidative damage, a function that likely evolved after the emergence of a well oxygenated post-GOE environment. Because oxygen is typically required for the synthesis of retinal, it is possible that earliest microbial rhodopsins could only have originated following the GOE or potentially the (likely hundreds of millions of years earlier) emergence of oxygenic photosynthesis. However, it has been demonstrated that rhodopsin-based phototrophy can also be carried out in hypoxic and anoxic conditions (DasSarma et al. 2012). In addition, an alternative, uncharacterized biosynthetic pathway for retinal synthesis has been reported (Nakajima et al. 2020; Chazan et al. 2022). Thus, the necessity of oxygen for retinal synthesis remains an open question. Despite ambiguity in the timing of rhodopsin origins relative to the GOE (Burnetti and Ratcliff 2020), it seems reasonable that the expansion of oxygenated niches following the GOE aided the ecological and functional diversification of rhodopsins, both expressed in the color tuning trends we observe in our phylogenetic analysis.

There are currently no absolute age constraints on either the oldest rhodopsin-bearing organisms or their biomolecular components, but an assessment of the distribution of light-driven sodium/proton export pumps among prokaryotes has been interpreted as indicating proximity of rhodopsin-like sodium translocation to the shared bacterial-archaeal superfamily ancestor (Shalaeva 2015). This would place the deepest proton-pumping ancestral nodes of our phylogenetic tree proximal to LUCA, with inferred ages of greater than ∼3.5 Ga, potentially preceding the divergence of the three domains of life (Ruiz-Gonzalez and Marin 2004). Considering such an early rhodopsin origin as a possible evolutionary scenario, one interpretation is that the earliest green-absorbing rhodopsins would have been well adapted to the water transmission peak at blue-green wavelengths in the marine early Earth surface environment – even at moderate depths in the ocean such that their host organisms could escape damaging UV rays prior to the accumulation of protective atmospheric ozone (Fig. 5a,b, Fig. 6). These spectral regions notably occupy the window excluded by the absorption peaks of chlorophylls and bacteriochlorophylls, potentially reflecting the evolutionary tradeoffs between retinal- and chlorophyll-/bacteriochlorophyll-based phototrophy (Burnetti and Ratcliff 2020)(Fig. 5d, Fig. 6b). Rhodopsin terrestrial and surface water host niches following the GOE might then have later been expanded by reduced UV irradiance, with shallow water niches more sharply discriminated by the increase in dissolved oxygen concentrations, contributing to the extant pattern of functional and sequence divergence of the rhodopsin protein family. Such a scenario would be broadly compatible with trends in mid-to-late Precambrian microfossil abundance, which (questions of taphonomic bias aside) consistently indicate significant microfossil diversity and abundance in near-shore marine deposits (Adam 2016; Javaux 2017). Host microorganisms residing in these photic zones would have been able to access available nutrients (e.g., phosphorous and nitrate), and avoid harmful UV radiation while simultaneously harnessing beneficial regions of the visible electromagnetic spectrum via the selective attenuation of light (and selective avoidance of dissolved oxygen) within the water column.

**Fig. 6.**
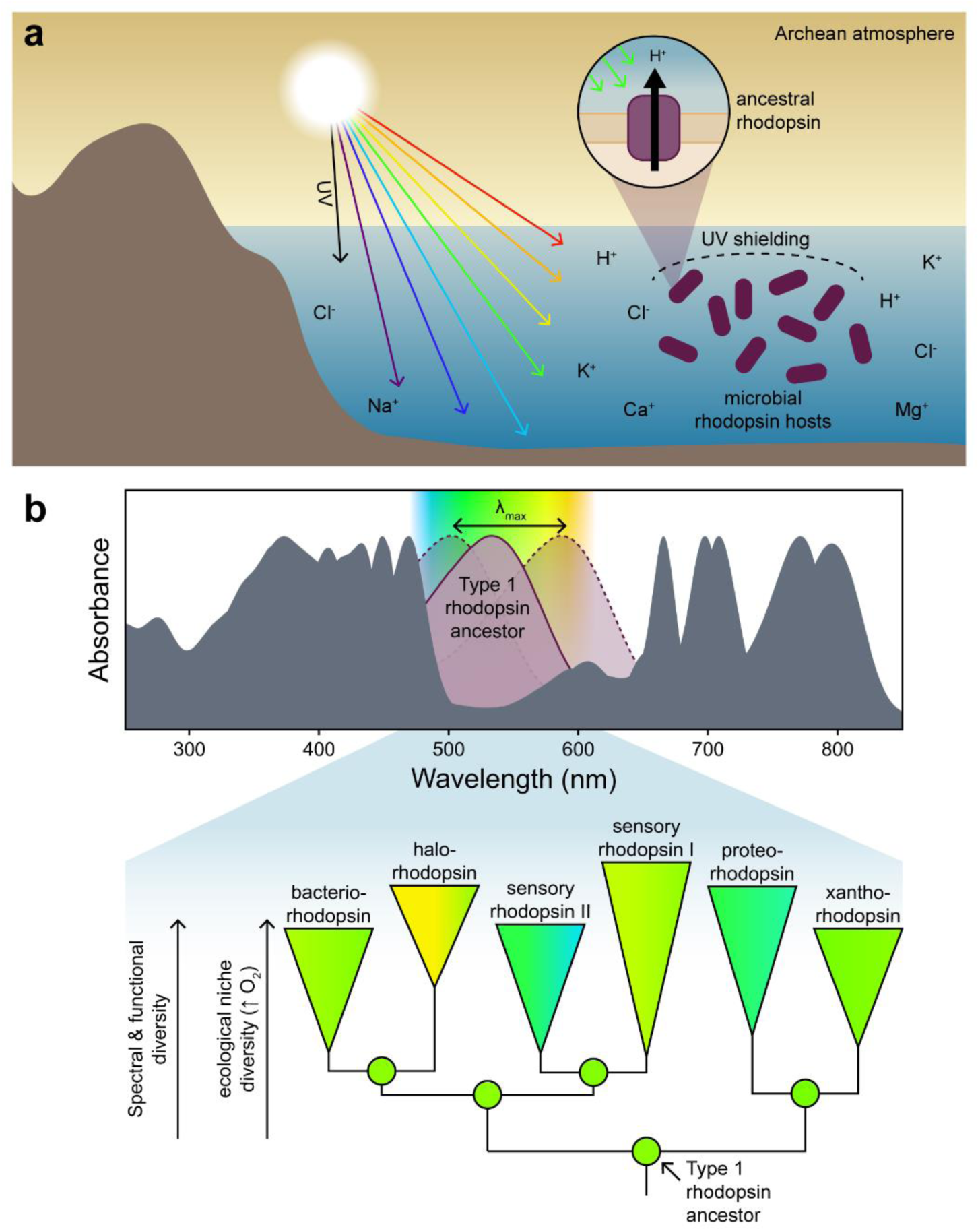
Coupling between surface irradiance, spectral tuning, and functional diversity over microbial rhodopsin evolution. (*A*) In a possible evolutionary scenario, microbial hosts of ancestral type 1 rhodopsins could have occupied depths in the Archean marine environment where high UV irradiation was attenuated and where they could have absorbed green wavelengths to which they were maximally tuned. These ancestral rhodopsins would likely have functioned as light-driven proton pumps, providing both a simplistic means of energy production and maintenance of the intracellular environment. (*B*) Color tuning of extant and ancestral microbial rhodopsins appear to explore the spectral window not already occupied by other biological pigments such as chlorophylls and bacteriochlorophylls (grey, adapted from Fig. 5d). Spectral diversification over rhodopsin evolutionary history likely accompanied functional specialization enabled by the expanded ecological niches following the rise of atmospheric oxygen.

### Implications for planetary habitability and the search for life beyond Earth

The coevolution of environment and life early in Earth’s history serves as a model for predicting universal, detectable biosignatures that might be generated on a microbe-dominated planet beyond our solar system. The current approach to constraining the possible suite of remotely detectable biosignatures leverages the aggregate spectral features of photosystems and pigment proteins which are sensitive to different portions of the Sun’s emission spectrum (Fig. 5) (Schwieterman et al. 2018). Photosystems, representing an early-evolved phototrophic strategy (Oliver et al. 2021), exhibit an immense capacity for tunability that is intimately associated with the diverse properties of attached pigments (Cardona 2015; Nürnberg et al. 2018). However, they are complicated, specialized structures, as opposed to rhodopsins which are comparatively simple and consolidate a photosensitive mechanism into a single protein. The dual roles of rhodopsin active site residues as determinants of both function and peak photon absorptivity demonstrate that these pigment proteins generate a suite of biosignatures that are tunable to our Sun’s peak irradiance profile, to long-term changes in the Earth surface chemical system, and to ecological niche diversification over billions of years. Our phylogenetic analysis indicates that rhodopsins’ structural simplicity, functional variability and spectral tunability make them an ideal testbed for assessing universal properties of biosignature production on candidate microbe-dominated planets that may differ from our referent biological (Earth-Sun) system.

A generic characteristic of life is that it is ‘tuned’ to the conditions found on our planet that enable its existence. These conditions refer to the chemical and physical characteristics that circumscribe habitability, namely substrate availability, ambient temperature, pH, atmospheric pressure and composition, salt concentration, and solar insolation, and irradiance, among other geochemical and atmospheric attributes. As demonstrated here, proteins can be far more tuned to these external conditions than other molecules or anatomical features that compose organisms, precisely because many proteins serve as intermediaries for channelizing photons and substrates into metabolic activity. Our results show that protein families (and to some extent, the niches occupied by their hosts) can be amenable to paleobiological reconstruction in ways that cannot be accomplished using conventional molecular biosignatures alone.

Utilizing proteins as paleosensors of geologic and stellar conditions in deep time, leveraging the informatic characteristics of protein composition, and the tight coupling between internal (cellular) and external (environmental) conditions, permits future studies exploring circumstances of habitability and biosignature production over Earth’s deep and varied history. Light absorbance profiles produced by microbial pigments can be used to prime future disk-integrated observations of exoplanets, provide surficial albedo parameters for complex atmospheric radiative transfer models, and directly aid the development of instrumentation for the detection and characterization of life in the universe both within and outside our Solar System. Forthcoming integrative studies that reconcile experimentally validated biological photon absorbances with more sophisticated planetary system analyses would thus be beneficial, for instance, in applying microbial spectral properties to assess the potential habitability or inhabitation of extrasolar planets (Hegde 2015; Schwieterman 2015; DasSarma 2020). Overall, understanding biomolecular adaptations may benefit planetary habitability assessments when planetary and stellar indicators are poorly constrained.

## METHODS

### Phylogenetic reconstruction and ancestral sequence inference

A type 1 microbial rhodopsin protein sequence dataset was created by identifying homologs from the National Center for Biotechnology Information non-redundant protein database in December 2020. Homologous sequences were identified and obtained via BLASTp (Camacho et al. 2009) using query amino acid sequences acquired from UniProt (Uniprot Consortium 2019) (Supplementary Table 6) and an expect value cutoff of <1e-5. The sequence dataset was curated to remove duplicate and partial sequences. The final dataset consisted of 568 sequences: 96 bacteriorhodopsin, 144 halorhodopsin, 83 proteorhodopsin, 29 sensory rhodopsin I, 56 sensory rhodopsin II, 44 xanthorhodopsin, and 116 outgroup heliorhodopsin sequences. The dataset was aligned by MAFFT (v7.427) (Katoh and Standley 2013) with a gap open penalty of 1.53 and offset value of 0.0. Fungal and algal type 1 rhodopsin sequences were excluded from the dataset due to long-branch-attraction effects observed within the bacterial rhodopsin clade. In addition, viral rhodopsin sequences (Yutin and Koonin 2012) were excluded due to the paucity of relevant paleobiological indicators for tracking their evolution in the context of Earth surface environmental changes.

The rhodopsin protein sequence alignment was used to select the best-fit evolutionary model by ModelFinder (Kalyaanamoorthy et al. 2017) implemented in the W-IQ-TREE web server(Trifinopoulos et al. 2016). Models accounting for the proportion of invariant sites (“+I”) were not included in model selection due to issues optimizing rate heterogeneity parameters independently (Yang 2006). The three top-ranking models scored by the Akaike and Bayesian Information Criterion, LG+G+F, WAG+G+F, and VT+G+F, were used for maximum likelihood phylogenetic inference by RAxML (v8) (Stamatakis 2014). Branch support was evaluated using 100 rapid bootstrap replicates. The type 1 rhodopsin clade was rooted with heliorhodopsin sequences, a protein group known to share a distant homology with type 1 rhodopsins (Pushkarev 2018; Shibukawa 2019). Because all tested evolutionary models yielded comparable topologies, the best-fit model, LG+G+F, was implemented for ancestral sequence inference by PAML (v4) (Yang 2007). Ancestral insertion/deletion sites were reconstructed using a binary likelihood model following Aadland et al. (2019).

### Homology modeling and functional predictions of ancestral rhodopsin sequences

Inferred maximum likelihood ancestral sequences for the five deepest phylogenetic nodes (Anc1-Anc5) were submitted to the Phyre2 web server (Kelley et al. 2015) for structural homology modeling. Selected structural templates for modeling and associated statistics are listed in Supplementary Table 7. Predicted ancestral rhodopsin structures were used for structure-based functional annotation by COFACTOR implemented in the I-TASSER software suite (Roy 2010), which predicts protein function based on global and local similarities to template proteins with known structures and functions.

### Alternative ancestral sequence library generation

For ancestral nodes Anc1-Anc5, 50 alternate protein sequences were generated from site-wise posterior probabilities to test the robustness of downstream analyses to statistical uncertainty of ancestral inference. For each ancestral node, regularly increasing percentages (>0% to 100%) of ambiguously reconstructed sites in the maximum likelihood sequence—defined here as having posterior probabilities <0.7—were replaced with the next most probable residue. Thus, the least probable ancestral sequence explored here by this approach had all ambiguous residues replaced by an alternate residue, likely representing the fringe of plausible ancestral sequence space (Eick et al. 2017).

### Wavelength Prediction

Ancestral maximum absorbance wavelength (λ_max_) values were predicted using the machine learning model, group-LASSO-based machine learning method (Karasuyama et al. 2018) implemented in R. This statistical model describes the relationship between primary amino acid sequence and rhodopsin color tuning based on a training rhodopsin sequence dataset with experimentally validated λ_max_ values. Maximum likelihood and alternate ancestral rhodopsin sequences, as well as all extant rhodopsin sequences used for phylogenetic inference, were aligned to the 796-sequence training dataset curated by Karasuyama et al. (2018) for λ_max_ prediction. To test the robustness of predicted λ_max_ to the size and composition of the training dataset, training data subsets were generated by 1) the sequence clustering method, CD-HIT (Fu et al. 2012) at 90%, 70%, or 50% sequence identity thresholds and 2) by randomly sampling 700, 500, 300, or 100 sequences from the original dataset. λ_max_ prediction for maximum likelihood rhodopsin sequences was repeated using the subsampled training datasets (Supplementary Table 5).

### Radiative transfer and pigment spectra

Surface photon irradiance spectra were calculated with the Spectral Mapping Atmospheric Radiative Transfer (SMART) model developed by D. Crisp (Meadows 1996). Calculation assumed cloud-free conditions with a solar zenith angle of 60 degrees and summed both direct and diffuse radiation at the surface. The modern Earth atmospheric profile was assumed to be a 1962 U.S. Standard Atmosphere employed by the InterComparison of Radiation Codes in Climate Models (ICRCCM) project (Luther 1988), whereas the Archean scenario was assumed to be a haze-free anoxic atmosphere from Arney et al. (2016) with a CH_4_/CO_2_ ratio of 0.1. The absorption coefficients of Segelstein (1981) and the Beer-Lambert Law were used to calculate the transmissivity and the attenuation of the photon irradiances as a function of depth.

## Data availability

All code and phylogenetic datasets can be found at https://github.com/kacarlab/Rhodopsin.

## ACKNOWLEDGEMENTS

We thank I. Takeuchi, M. Karasuyama, and K. Inoue for access to the machine learning algorithms used to predict rhodopsin wavelength absorbances used in this work, for the supportive assistance and the useful comments, and S. DasSarma for the insightful conversations. This work was supported by the NASA Arizona Space Grant (C.D.S.), the NASA Astrobiology Postdoctoral Fellowship, (A.K.G.), the NASA Interdisciplinary Astrobiology Research Consortium (ICAR) MUSE, a member of the NASA Early Cell to Multicellularity Research Coordination Network funded via the Astrobiology Research Program No 80NSSC21K0592 (B.K.), the NASA Early Career Faculty Award No. 80NSSC19K1617 (B.K.) and an NSF Emerging Frontiers Program Award No. 1724090 (B.K.), NASA’s Alternative Earths team funded by the NASA Astrobiology Institute under Cooperative Agreement Number NNA15BB03A and the NASA ICAR program, and the Virtual Planetary Laboratory, a member of the NASA Nexus for Exoplanet System Science (NExSS) funded via NASA Astrobiology Program Grant No. 80NSSC18K0829 (E.W.S.).

## AUTHOR CONTRIBUTIONS

B.K. conceived, designed, and supervised the overall study. C.D.S. designed and performed the phylogenetic and modeling experiments. E.F. designed and generated the ancient protein library, wavelength prediction and robustness analysis pipeline with input from A.K.G. E.W.S. conducted the radiative transfer calculations and contributed to the habitability at depth hypothesis with input from Z.R.A. All authors discussed and interpreted the results. B.K. and C.D.S. wrote the manuscript with input and/or text contributions from all authors.

