## Supplementary Information for "Earliest photic zone niches probed by ancestral microbial rhodopsins"

### Earliest photic zone niches probed by reconstruction of ancestral bacterial rhodopsin proteins

#### SUPPLEMENTARY INFORMATION

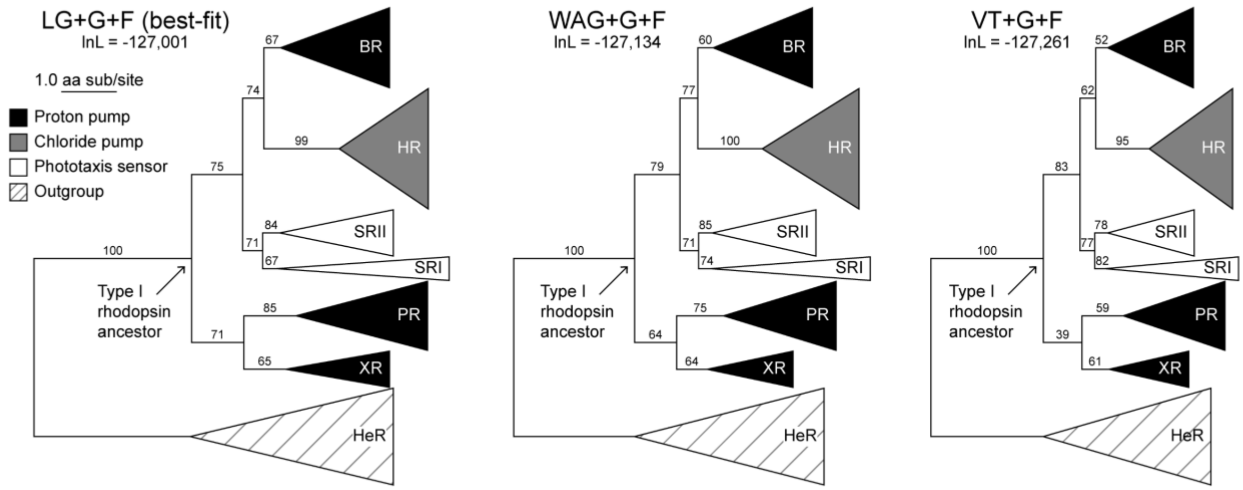


**Supplementary Fig. 1 Robustness of microbial type I rhodopsin protein phylogeny to three of the best-fitting evolutionary models.** Log likelihood (“lnL”) scores decrease from left to right. Branch support values are derived from 100 bootstrap replicates. Branch scale represents 1.0 amino acid substitutions per site. Legend applies to all trees. BR: bacteriorhodopsin; HR, halorhodopsin; SR II, sensory rhodopsin II; SR I, sensory rhodopsin I; PR, proteorhodopsin; XR, xanthorhodopsin; HeR, heliorhodopsin.

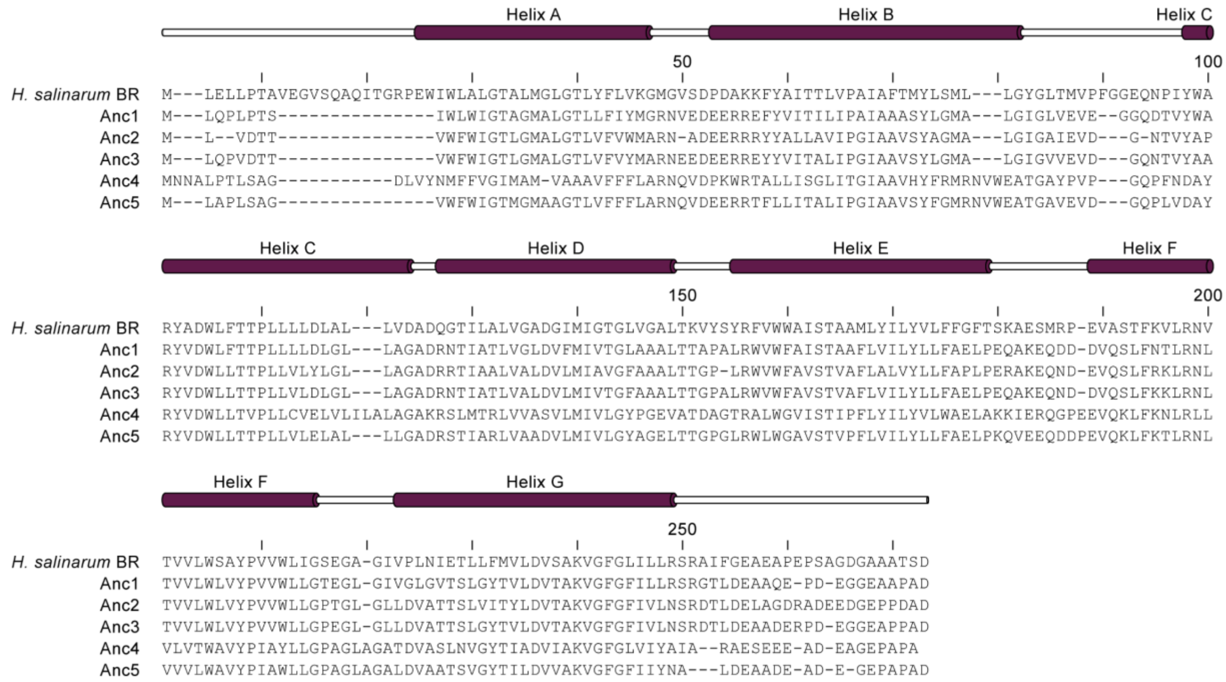


**Supplementary Fig. 2 Protein sequence alignment of extant (*Halobacterium salinarum* bacteriorhodopsin, "BR") and ancestral rhodopsins.** Transmembrane helix locations shown above sequence alignment.

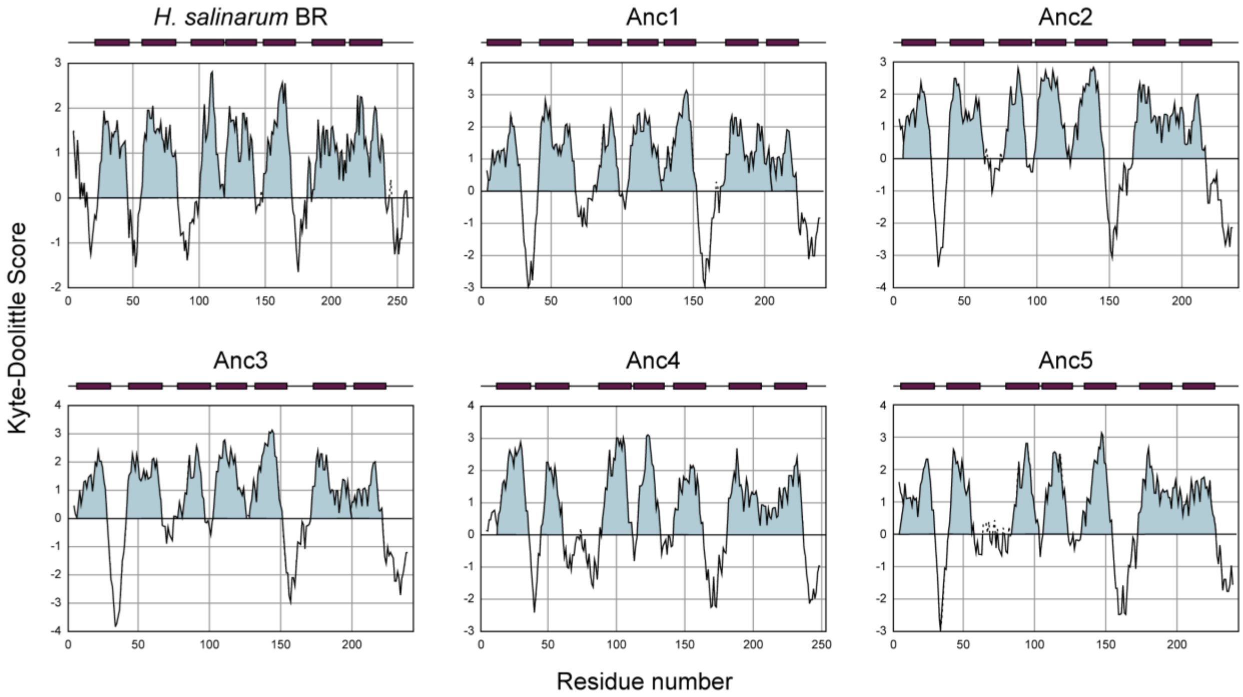

**Supplementary Fig. 3 Hydropathy plots for extant (*Halobacterium salinarum* bacteriorhodopsin, "BR") and ancestral rhodopsins.** Hydrophobic domains (Kyte-Doolittle score > 0) are shaded and approximate positions of transmembrane helices are indicated by purple bars above each plot.

**Supplementary Table 1 Mean site-wise posterior probabilities of ancestral rhodopsins, calculated for full sequence length or for 23 binding pocket residues.**

| **Ancestor** | **Mean posterior probability, full sequence (± 1σ)** | **Mean posterior probability, binding pocket (± 1σ)** |
|---|---|---|
| Anc1 | 0.76 (± 0.24) | 0.96 (± 0.11) |
| Anc2 | 0.76 (0.25) | 0.95 (± 0.14) |
| Anc3 | 0.71 (± 0.26) | 0.95 (± 0.12) |
| Anc4 | 0.65 (0.28) | 0.97 (± 0.06) |
| Anc5 | 0.55 (± 0.29) | 0.90 (± 0.16) |

**Supplementary Table 2 Site-wise posterior probabilities of ancestral rhodopsin binding pocket residues, homologous to the listed residues in extant *Halobacterium salinarum* bacteriorhodopsin.**

| **Homologous residue in extant BR** | **Posterior probability** | | | | |
|---|---|---|---|---|---|
| | **Anc1** | **Anc2** | **Anc3** | **Anc4** | **Anc5** |
| Y83 | 1.00 | 1.00 | 1.00 | 1.00 | 0.996 |
| D85 | 1.00 | 1.00 | 1.00 | 1.00 | 0.997 |
| W86 | 1.00 | 1.00 | 1.00 | 1.00 | 0.998 |
| T89 | 0.995 | 1.00 | 1.00 | 1.00 | 0.999 |
| T90 | 1.00 | 1.00 | 1.00 | 0.977 | 0.836 |
| P91 | 1.00 | 1.00 | 1.00 | 1.00 | 1.00 |
| L93 | 0.989 | 0.997 | 0.990 | 0.719 | 0.759 |
| D96 | 0.711 | 0.945 | 0.540 | 0.984 | 0.605 |
| M118 | 1.00 | 0.999 | 1.00 | 1.00 | 1.00 |
| I119 | 0.994 | 1.00 | 0.999 | 0.947 | 0.979 |
| G122 | 1.00 | 1.00 | 1.00 | 1.00 | 1.00 |

| | | | | | |
|---|---|---|---|---|---|
| W138 | 0.608 | 0.994 | 0.746 | 0.943 | 0.493 |
| S141 | 1.00 | 0.995 | 0.999 | 0.999 | 0.992 |
| T142 | 0.785 | 0.521 | 0.725 | 0.943 | 0.810 |
| M145 | 1.00 | 1.00 | 1.00 | 0.887 | 0.909 |
| W182 | 1.00 | 1.00 | 1.00 | 1.00 | 0.999 |
| Y185 | 1.00 | 1.00 | 1.00 | 1.00 | 0.995 |
| P186 | 1.00 | 1.00 | 1.00 | 1.00 | 1.00 |
| W189 | 1.00 | 1.00 | 0.999 | 0.992 | 0.517 |
| F208 | 0.971 | 0.517 | 0.970 | 1.00 | 0.995 |
| D212 | 1.00 | 1.00 | 1.00 | 1.00 | 0.999 |
| A215 | 0.999 | 0.979 | 0.984 | 0.888 | 0.900 |
| K216 | 1.00 | 1.00 | 1.00 | 1.00 | 1.00 |

**Supplementary Table 3 SOSUI[1] predictions for subcellular localization and hydrophobicity of ancestral rhodopsins.**

| Ancestor | Protein localization | Average Hydrophobicity score |
|---|---|---|
| Anc1 | Membrane | 0.670 |
| Anc2 | Membrane | 0.644 |
| Anc3 | Membrane | 0.562 |
| Anc4 | Membrane | 0.655 |
| Anc5 | Membrane | 0.668 |

**Supplementary Table 4 I-TASSER COFACTOR[2] structure-based function annotations for ancestral rhodopsins.**

| Ancestor | Biological Process (GO Term \| probability) | Cellular Component (GO Term \| probability) | Molecular Function (GO Term \| probability) |
|---|---|---|---|
| Anc1 | Phototransduction (GO: 0007602 \| 0.990) | Integral component of Membrane (GO: 0016021 \| 1.000, 0.990) | Ion channel activity (GO: 0005216 \| 1.000) |

| | | | |
|---|---|---|---|
| | Protein – chromophore linkage<br>(GO: 0018298 \| 0.990)<br><br>Proton transmembrane transport<br>(GO: 0015992 \| 0.970) | Plasma membrane<br>(GO: 0005886 \| 0.909, 0.990) | Photoreceptor activity<br>(GO: 0009881 \| 0.990) |
| Anc2 | Protein-chromophore linkage<br>(GO: 0018298 \| 0.990)<br><br>Phototransduction<br>(GO:0007602 \| 0.990)<br><br>Proton transmembrane transport<br>(GO: 0015992 \| 0.970) | Integral component of membrane<br>(GO:0016021 \| 1.000, 0.99)<br><br>Plasma membrane<br>(GO:0005886 \| 0.916, 0.990) | Ion channel activity<br>(GO:0005216 \| 0.990)<br><br>Photoreceptor activity<br>(GO:0009881 \| 0.990) |
| Anc3 | Phototransduction<br>(GO: 0007602 \| 0.990)<br><br>Protein – chromophore linkage<br>(GO: 0018298 \| 0.990)<br><br>Proton transmembrane transport<br>(GO: 0015992 \| 0.970) | Integral component of Membrane<br>(GO: 0016021 \| 1.000, 0.990)<br><br>Plasma membrane<br>(GO: 0005886 \| 0.922, 0.990) | Ion channel activity<br>(GO: 0005216 \| 1.000)<br><br>Photoreceptor activity<br>(GO: 0009881 \| 0.990) |
| Anc4 | Protein-chromophore linkage<br>(GO: 0018298 \| 0.980)<br><br>Phototransduction<br>(GO:0007602 \| 0.980) | Integral component of membrane<br>(GO:0016021 \| 1.000, 0.98)<br><br>Plasma membrane | Ion channel activity<br>(GO:0005216 \| 0.980)<br><br>Photoreceptor activity |

| | | | |
|---|---|---|---|
| | Proton transmembrane transport<br>(GO: 0015992 \| 0.960) | (GO:0005886 \| 0.980) | GO:0009881 \| 0.980 |
| Anc5 | Phototransduction<br>(GO: 0007602 \| 1.000)<br><br>Protein – chromophore linkage<br>(GO: 0018298 \| 1.000)<br><br>Proton transmembrane transport<br>(GO: 0015992 \| 0.990) | Integral component of Membrane<br>(GO: 0016021 \| 1.000, 1.000)<br><br>Plasma membrane<br>(GO: 0005886 \| 1.000) | Ion channel activity<br>(GO: 0005216 \| 1.000)<br><br>Photoreceptor activity<br>(GO: 0009881 \| 1.000) |

**Supplementary Table 5 Robustness of the group-LASSO-based maximum absorbance wavelength ($\lambda_{max}$) prediction method[3] to size and composition of the training dataset (originally 796 sequences), subsampled by sequence clustering (by CD-HIT[4]) or random selection.**

| Subsampling method | Training dataset size (# of sequences) | Predicted $\lambda_{max}$ | | | | |
|---|---|---|---|---|---|---|
| | | **Anc1** | **Anc2** | **Anc3** | **Anc4** | **Anc5** |
| No subsampling | 796 | 537 | 540 | 542 | 543 | 541 |
| CD-HIT, 90%* | 73 | 538 | 538 | 538 | 538 | 538 |
| CD-HIT, 70%* | 54 | 541 | 541 | 541 | 541 | 541 |
| CD-HIT, 50%* | 34 | 530 | 529 | 528 | 529 | 530 |
| Random | 700 | 534 | 539 | 540 | 541 | 539 |
| Random | 500 | 532 | 542 | 540 | 541 | 540 |
| Random | 300 | 524 | 537 | 539 | 537 | 536 |
| Random | 100 | 540 | 539 | 551 | 556 | 546 |

*Sequence identity threshold for CD-HIT clustering

**Supplementary Table 6 BLAST query sequences used for extant rhodopsin homolog identification.**

| Rhodopsin Type | Accession Number |
|---|---|
| Bacteriorhodopsin | P02945 (BACR_HALSA) |
| Halorhodopsin | B0R2U4 (BACH_HALS3) |
| Sensory rhodopsin I | P0DMH8 (BACS1_HALSA) |
| Sensory rhodopsin II | P71411 (BACS2_HALSA) |
| Blue proteorhodopsin | Q9AFF7 (PRRB_PRB02) |
| Green proteorhodopsin | Q9F7P4 (PRRG_PRB01) |
| Xanthorhodopsin | Q2S2F8 (Q2S2F8_SALRD) |

**Supplementary Table 7 Phyre2[5] protein structure prediction statistics for ancestral rhodopsin sequences.**

| Ancestor | Phyre Template ID (PDB ID) | Confidence | % ID | Coverage | Type (Organism) |
|---|---|---|---|---|---|
| Anc1 | C4jr8A (4JR8) | 100% | 61% | 92% | Cruxrhodopsin-3 (*Haloarcula vallismortis*) |
| Anc2 | C4fbzA (4FBZ) | 100% | 50% | 93% | Deltarhodopsin (*Haloterrigena thermotolerans*) |
| Anc3 | C4fbzA (4FBZ) | 100% | 52% | 94% | Deltarhodopsin (*Haloterrigena thermotolerans*) |
| Anc4 | C3ddlB (3DDL) | 100% | 55% | 95% | Xanthorhodopsin (*Salinibacter ruber*) |
| Anc5 | C3ddlB (3DDL) | 100% | 42% | 95% | Xanthorhodopsin (*Salinibacter ruber*) |
